# Distribution of the National Science Foundation’s Advancing Informal STEM Learning Awards (AISL) between 2006-21

**DOI:** 10.1101/2022.02.18.480415

**Authors:** Heidi M. Houzenga, Fanuel J. Muindi

**Author notes:** (Corresponding author)* Fanuel J. Muindi. **Contact Information** Heidi Houzenga.

## Abstract

The COVID-19 pandemic tested many fundamental connections between science and society. A growing field working to strengthen those connections exists within the informal STEM learning (ISL) community which provides diverse learning and engagement environments outside the formal classroom. One of the largest funders of ISL initiatives is the National Science Foundation (NSF) which runs the Advanced Informal STEM Learning (AISL) program in the United States. The AISL program supports initiatives through six categories that include pilots and feasibility studies, research in service to practice, innovations in development, broad implementation, literature reviews, syntheses, or meta-analyses, and conferences. However, a number of questions remain unanswered with respect to the distribution and ultimately the broad impact of the awards. In this study, we analyzed publicly available awardee information across a 15-year period (2006-2021) to provide a preliminary analysis of how the AISL grants are distributed across individual states, organizations, and the principal investigators. Several states, organizations, and principal investigators stood out as prolific awardees. Massachusetts and California represented the largest share of awards at 14% and 13% respectively during that time. WGBH Educational Foundation located in Massachusetts and the Exploratorium located in California, received the largest number of awards during the 15 year period. Notably, 67% of the AISL awards list at least one co-principal investigator. Our report brings to light a number of new questions and charts new paths of exploration for future studies.

## INTRODUCTION

Informal learning experiences in science, technology, engineering, and math (STEM) involve diverse settings outside of the formal classroom. This includes museums, zoos, science festivals, social media, blogs, after school programs, and many other local community science events. There is a growing body of work that show informal STEM learning (ISL) initiatives provide an important supplementation of the formal school experience and ultimately help improve students’ awareness of and interest in the STEM fields (Denson et al., 2015; Mohr-Schroeder et al., 2014; Roberts et al., 2018).

The National Science Foundation (NSF) is a federal agency tasked with supporting the field of science and engineering and is one of the largest funders of ISL initiatives in the world through its Advancing Informal STEM Learning AISL (AISL) program. At the core, the AISL program seeks to advance the design and engagement of ISL opportunities for the public and expand research on and assessment of ISL environments (James & Singer, 2016). In the last several years, the AISL program has been funding programs within the following six categories: pilots and feasibility studies; research in service to practice; innovations in development; broad implementation; literature reviews; syntheses, or meta-analyses; and conferences.

As a significant funder of ISL initiatives, the NSF has a significant influence with respect to the growth of the field. Surprisingly, there are a number of questions that are not well analyzed. For example, what type of initiatives and research have been funded? How has this changed over time? Who is receiving these awards? What is the distribution and size of these awards among states and host organizations? These and many other questions are important to know and track over time as they provide an insight into the direction the field is going since this is where many of the top ideas in ISL come for large scale funding. As such, the annual funding competition is extremely competitive. However, even though much of the awardee information is public (the NSF does provide users with a searchable database), the output from the database has yet to be deeply analyzed, visualized, and discussed publicly to our knowledge. In this brief report, we provide the first attempt to answer some of the aforementioned questions and discuss some paths for future studies.

## APPROACH

All data represented was obtained through the publicly made available National Science Foundation online award database (https://nsf.gov/awardsearch/advancedSearch.jsp). The advanced search feature was utilized with element code 7259 which represents the Advancing Informal STEM Learning awards. Active and expired awards within a 15 year period between 09/01/2006 to 09/01/2021 were exported to a CSV file for analysis. In total, 1,043 award data points were reported. Categories of interest in the analysis included dates, principal investigators, states, organizations, and co-principal investigators. During the 15 year period, 764 principal investigators, 48 states (including the District of Columbia and Puerto Rico), and 393 organizations were recipients of AISL awards. GraphPad Prism version 9.0.0 for Windows/Mac, GraphPad Software, San Diego, California USA, www.graphpad.com, was used to visualize the results.

## RESULTS

### AISL Grants across States

Addressing the question of what states are receiving awards, we find 48 states and territories are represented in the 1,043 awards issued in the 15 years. The overall state information is shown in Figure 1. Figure 1A reflects the number of states to the number of awards issued in those 15 years. Several states of interest stand out as receiving 20 or more awards (Figure 1B). Massachusetts (Figure 1C) comes in as the leader in states with 144 awards, reflecting 14% of the total in the 15 year period, and has consistently received at least four awards every year. The state of California (Figure 1D) is also a prolific awardee coming in with 137 awards or 13% of the total awards. Like Massachusetts, they have also consistently received an award in every year of the 15 years. Other states from Figure 1B that have received more than 20 awards during the time include New York, Illinois, District of Columbia, Pennsylvania, Washington, Colorado, Oregon, Minnesota, Maryland, North Carolina, Virginia, Arizona, and Florida. 25 states and territories had received 1-10 awards. Missing within the 15 year period are the states of Alabama, Arkansas, Delaware, and South Dakota. No AISL awards have been issued to those states. Due to lack of data, it remains unknown whether the states with no AISL awards are being unsuccessful with their applications or they are simply not applying all together. However, it is worth mentioning that many of the grants are not confined to within a specific state and as such, other states do end up benefiting as well.

**Figure 1:**
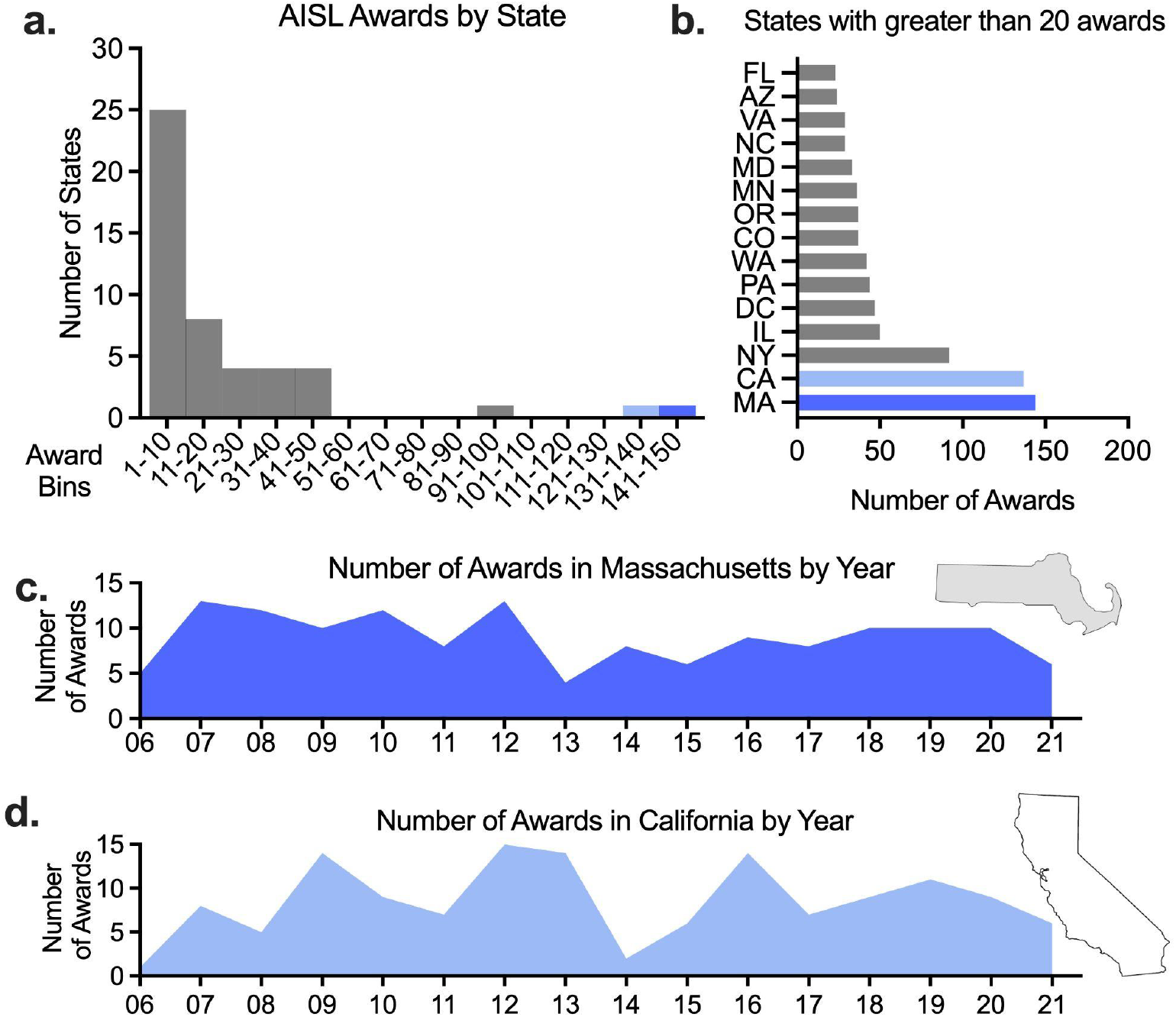
AISL Grants across States. **(A)**The number of awards received per state for the years 2006-2021. The number of awards (x-axis) is placed in bin widths of 10 awards. **(B)** States that have received more than 20 awards in the years 2006-2021. Dark blue represents the state with the most awards in those years and light blue, the state with the second-most awards. **(C)** The number of awards received per year by the state of Massachusetts for the years 2006-2021. **(D)** The number of awards received per year by the state of California for the years 2006-2021.

### AISL Grants across Organizations

Organization data for AISL awards is shown in Figure 2. 393 organizations have acquired awards during this 15 year period. While the majority of organizations seen in Figure 2A have one to five awards, we again see several organizations that stand out. Six organizations are highlighted in Figure 2B as receiving more than 15 AISL awards. WGBH Educational Foundations (Figure 2C), located in Massachusetts, has the most awards in the past 15 years with 31. They have received awards in 13 out of the past 15 years. Exploratorium (Figure 2D), located in California, was awarded 23 times and they have also received awards in 13 out of the 15 years. TERC, Inc, located in Massachusetts, and the University of Washington both received 21 awards. The Museum of Science, located in Massachusetts, and Northwestern University, located in Illinois, have received 16 and 15 awards respectively. 205 out of the 393 are organizations that received a single award.

**Figure 2:**
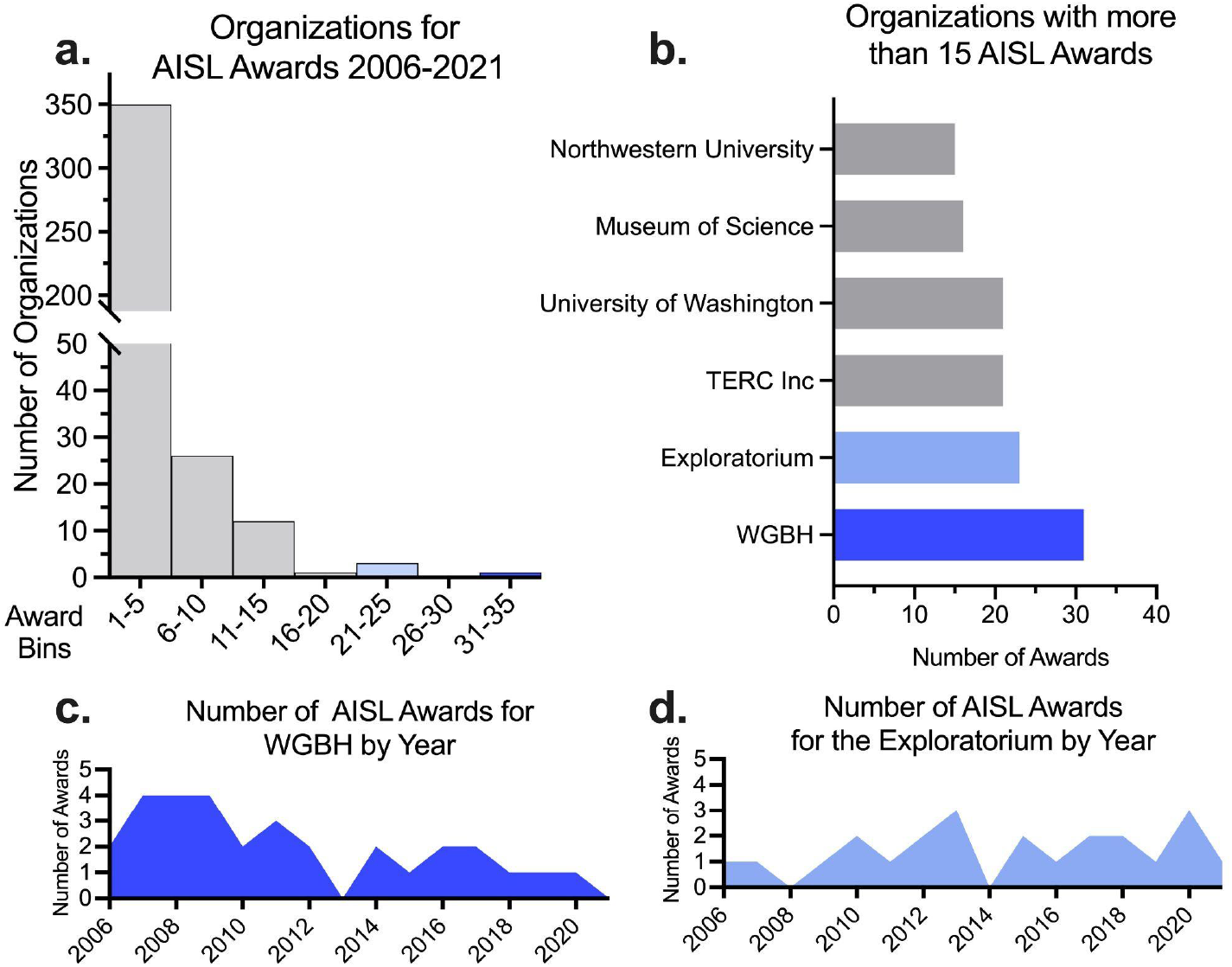
AISL Grants across Organizations. **(A)** The number of awards received by organizations for the years 2006-2021. The number of awards (x-axis) is placed in bin widths of 5. **(B)** Organizations that have received more than 15 awards in the years 2006-2021. Dark blue represents the organization with the most awards in those years and light blue, the organization with the second most awards. **(C)** The number of awards received per year by WGBH Educational Foundation for the years 2006-2021. **(D)** The number of awards received per year by Exploratorium for the years 2006-2021.

### AISL Grants across Principal Investigators

Analysis of principal investigators is shown in Figure 3. Although the majority of principal investigators have received a single award, we see in Figure 3A that several principal investigators have had great success in receiving multiple awards over these years. Marisa Wolsky, the principal investigator for WGBH, has received nine awards in the 15 year time period, most of those with a co-principal investigator. Kate Taylor, also with WGBH, received 8 awards, with three awards with co-principal investigator Marisa Wolsky. The analysis of the NSF database to assess the level of awards with more than one principal investigator (Figure 3B) reveals 700 out of the 1043 awards or 67% of the AISL awards in the 15 years have a co-principal investigator.

**Figure 3:**
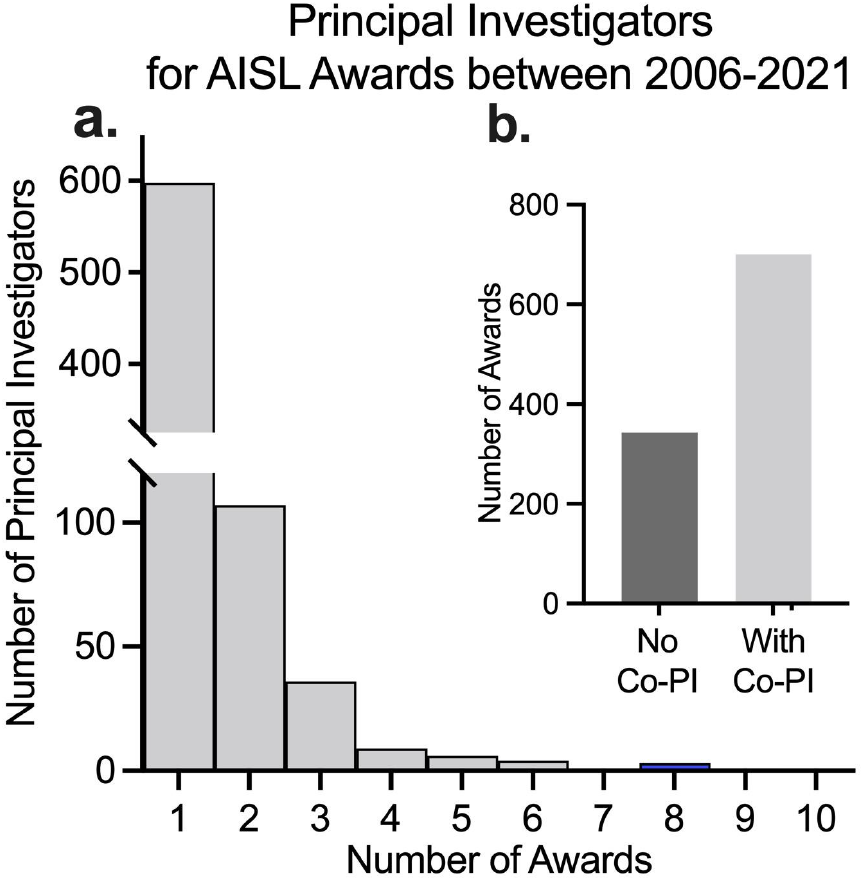
AISL Grants across Principal Investigators. **(A)** The number of awards received by principal investigators for the years 2006 to 2021. **(B)** Awards listed with or without a co-principal investigator (Co-PI).

## DISCUSSION

### The missing links

Overall, the results here provide an overview of the distribution of AISL grants across individual states, organizations, and the principal investigators across a 15-year period. The results bring to focus a number of new questions. For example, who is applying to the NSF AISL awards? Due to lack of data, it remains unknown whether the states with no AISL awards are being unsuccessful with their applications or they are simply not applying all together. No matter the situation, knowing this information will ultimately help the NSF decide how to target their outreach and education for organizations with respect to their size, location, and target audiences. Another question pertains to the organizations and the investigators that are being successful across years. Specifically, what is enabling their success? Such a question would require a close examination of those organizations and what the level of support they provide their investigators to enable them to successfully apply and eventually win such grants. At the same time, getting insights from those that were rejected will also be just as important, if not more, especially those that get reviewer comments.

The questions surrounding impact were not addressed in this report. A key question here entails comparing the levels of impact that the funded awards had on their intended audiences. The growing repository of publicly available reports (https://www.informalscience.org) from previous awards provides a rich data set for deep analysis into this question. A critical challenge there is coming up with a standard set of metrics to allow comparison across awards within a given funding category. Such an assessment will likely yield useful insights into the areas where the funding is having a large impact, where it is not, and ultimately help decide on how to best invest available funding in those respective areas moving forward. Another question surrounding impact pertains to the type of outcome(s) involved (awareness, knowledge, understanding, attitudes, skills, behavior, etc.) (Friedman, 2008) against the AISL program’s priorities of maximizing strategic impact, enhancing knowledge-building, promoting innovation, advancing collaboration, strengthening infrastructure, and broadening participation. Can the key outcome of one funded award result in a higher ranking with respect to the overall impact relative to another grant with a different key outcome within a specific priority area?

### Capture the human stories

Amidst the analysis, it is very easy to forget that at the end of the day, human beings are applying to these awards and doing the work. Their stories matter. As such, collecting and analyzing these stories will provide unique insights for all stakeholders involved. For example, such an analysis would provide context into what the applicants are facing within their organizations in both applying and also waiting for the news that comes many months later. These stories are in fact even more critical for those applicants that are not selected. How are they affected by not getting funded? Do they reapply? What support do they wish they had during the application process? What is the impact on their projects? For those funded, what additional support do they wish they received during the funding period? How did they manage their collaborations? These are just some of the many questions one could discover by going beyond the numbers and capturing the human stories orbiting the AISL program.

### Collaboration matters

As noted in Figure 3, we discovered that a large number of proposals have a co-PI. This was not a huge surprise given that one of AISL’s priority areas is advancing collaboration thus we suspect some number of proposals target that priority area. In fact, a research and evaluation company Rockman et al recently noticed that the depth and authenticity of collaborations across organizations is something that reviewers are paying close attention to (“AISL Review Criteria,” 2021). Two noteworthy questions they noted for potential co-PIs to consider are (1) the representation of community demographics, knowledge and values amongst the core team and (2) how the collaboration will ensure that all voices are represented equitably. Ultimately, connecting how a specific collaboration will address one or more of AISL’s priority areas will be critical.

### Breaking down the categories

It is also worth mentioning that although the NSF Merit Review provides overall demographics and funding rates for proposals (*NSF Merit Review Process Fiscal Year Digest*, 2019), those specific details for AISL awards were not publicly available. The NSF awards advanced search provides AISL award information. What could not be found in the search features are demographics, funding rates, and a way to decipher the six project types that AISL supports across the 15-year period. This is important because the AISL program supports six different types of awards which include Pilots and Feasibility Studies, Research in Service to Practice, Innovations in Development, Broad Implementation, Literature Reviews, Syntheses, or Meta-analyses, and Conferences (*Advancing Informal STEM Learning (AISL) (Nsf21599)* | *NSF - National Science Foundation*, n.d.). Future research direction could entail specific demographics and funding rates specific to AISL awards and the six supporting program categories. The breakdown of the funded awards by the different project types in addition to the audiences being targeted will be critical to get higher resolution with what is being funded and who the funding is impacting.

### Final Words

The analysis here provides an initial glance into the distribution of the AISL awards across states, organizations, and principal investigators. It is designed to serve as a catalyst for generating new questions and discussions. The much deeper analysis is for future studies to dig further into. The creation of a public dashboard using platforms such as Tableau could make it easier for the public, practitioners, and researchers to get a visual summary of all AISL awards and better digest the information across multiple dimensions. Undoubtedly, AISL awards will continue to serve as a powerful engine of innovation in informal STEM learning space in the decades to come and help ensure strong connections between science and society that are tough enough to withstand the stress of both ongoing and future societal challenges.

## ACKNOWLEDGEMENTS

We would like to thank Jessica W. Tsai and Sarah Dunifon for their feedback and all the members of the STEM Advocacy Institute (SAi) for their support.

## AUTHOR CONTRIBUTION STATEMENT

H.M.H. and F.J.M. jointly conceived the idea for the research project. H.M.H performed the data organization and subsequent analysis with supervision and verification from F.J.M. Both authors discussed the results and contributed to the final manuscript.

